# Implementation of the Water Framework Directive: Lessons learned and future perspectives for an ecologically meaningful classification of the status of Greek lakes, Mediterranean region

**DOI:** 10.1101/371799

**Authors:** Maria Moustaka-Gouni, Ulrich Sommer, Athena Economou-Amilli, George B. Arhonditsis, Matina Katsiapi, Eva Papastergiadou, Konstantinos A. Kormas, Elisabeth Vardaka, Hera Karayanni, Theodoti Papadimitriou

## Abstract

The enactment of the Water Framework Directive (WFD) initiated scientific efforts to develop reliable methods for comparing prevailing lake conditions against *reference* (or non-impaired) states, using the state of a set biological elements. Drawing a distinction between impaired and natural conditions can be a challenging exercise, as it stipulates the robust delineation of reference conditions along with the establishment of threshold values for key environmental variables used as proxies for the degree of system impairment. Another important aspect is to ensure that water quality assessment is comparable among the different Member States. In this context, the present paper offers a constructive critique of the practices followed during the WFD implementation in Greece by pinpointing methodological weaknesses and knowledge gaps that undermine our ability to classify the ecological status of Greek lakes. One of the pillars of WDF is a valid lake typology that sets ecological standards transcending geographic regions and national boundaries. The national typology of Greek lakes has failed to take into account essential components (e.g. surface area, altitude, salinity). WFD compliance assessments based on descriptions of phytoplankton communities are oversimplified and as such should be revisited. Exclusion of most chroococcal species from the analysis of cyanobacteria biovolume in Greek lakes and most reservoirs in the Mediterranean Geographical Intercalibration Group (Greece, Spain, Portugal and Cyprus) is not consistent with the distribution of those taxa in lakes. Similarly, the total biovolume reference values and the indices used in their classification schemes reflect misunderstandings of WFD core principles. This hampers the comparability of ecological status across Europe and leads to quality standards that are too relaxed to provide an efficient target especially for the protection and management of Greek/transboundary lakes such as Lake Megali Prespa, one of the oldest lakes in Europe.

## INTRODUCTION

The WFD (European Commission 2000) is considered the most important and innovative European environmental legislation for maintaining and improving the aquatic environment in the European Union (Boeuf and Fritsch 2016). The main objective of the WFD is that all surface waters should be in good or better ecological status by 2015 or at the latest in 2027. To achieve the “good status” objective, the Member States (MSs) should define and implement the necessary remediation programs. WFD is the first legislation establishing the innovative concept of ecological status of surface waters based on Biological Quality Elements (i.e. phytoplankton) in addition to physical and chemical conditions used in traditional legislation. As articulated in Directive definitions (Article 2), ecological status is an expression of the quality of the structure and functioning of aquatic ecosystems. MSs have the mandate to develop assessment methods of the ecological status for all biological quality elements listed in the normative definitions given in the Directive’s Annex V (European Commission 2000).

The ecological quality assessment based on ecological quality ratios (EQR; the ratio of observed to reference values) requires the delineation of type-specific reference conditions, as an essential piece of knowledge for distinguishing between “good” and “moderate” ecological conditions. Climatic and geological variability prohibits the establishment of absolute standards for the entire Community, and therefore type-specific biological reference conditions should be defined by MSs as targeted standards for maintaining and restoring ecological status. To ensure comparable definitions of good ecological status across Europe, MSs are also responsible for inter-calibrating the good ecological status class boundaries of their methods for each biological element in each water category with other MSs having common types of water bodies. Therefore, scientifically sound types design has a key role in setting standards to adequately partition prevailing ecological variation (i.e. in phytoplankton and macrophytes).

Lakes are usually assigned to types according to morphometric, geological, and altitudinal properties. Each MS can develop its own typology in System B by using common descriptors interpreted differently (i.e. different lake mean depths as thresholds for “deep lakes”) or different descriptors. Nevertheless, the common types do not include the lake surface area as an operational descriptor although it is prescribed as an obligatory descriptor by WFD, System A (ANNEX II) (European Commission 2000). Despite the intention of WFD to establish comparable typology standards across the Community, the existing practices have resulted in diverse national standards, even though the underlying ecological mechanisms are not geographically diverse and aquatic ecology has no national boundaries (Moss 2008). The typology descriptors should ensure that the derivation of type-specific biological reference conditions is defensible and thus the classification of lake ecological status, the heart of WFD, is reliable. Following this line of reasoning, it can be inferred that crucial to any implementation of the WFD is to first establish what reference (natural or undisturbed) conditions mean in terms of ecological variables suitable for monitoring programs (e.g. biomass, abundance, taxonomic composition, functional traits) based on sound limnological principles.

Defining biological reference conditions and setting boundaries for ecological status classes continues to represent a major challenge across Europe, and thus effective guidance by the scientific community appears to be a decisive factor for operational implementation of the framework. The definition should be based on transparent analyses of empirical data from a network of references sites if available. Based on new findings, Baattrup-Pedersen et al. (2017) suggest that more effort should be directed at describing reference conditions by experts instead of focusing solely on the development of assessment systems using pressure-impact frameworks. Each GIG or even MSs have their own diagnostics for identifying reference sites, occasionally with insufficient clarification of the methods chosen by authorities and insufficient clarification of legislative classification of the water bodies (e.g. Stoyneva et al. 2014).

For ecological status classification, hundreds of indices characterizing the status of biological elements have been developed during WFD implementation. Most indices have limitations for a variety of reasons, including applicability within a limited biogeographical area; reliance on a list of taxa rather than structure and functions; correlation rather than causation; high precision but poor accuracy; and poorly defined reference conditions (e.g. Moss 2008). Several indices are complex and expensive and therefore unsuitable to address the need for cost-effective and scientifically defensive ecological assessment (e.g. Katsiapi et al. 2016; Baattrup-Pedersen et al. 2013, 2017). Ecological quality of lakes judged by experts can be predicted reasonably well and affordably from water transparency, expressed as Secchi Disc depth (Peeters et al. 2008; Katsiapi et al. 2016). Among biological indicators for water quality assessment, phytoplankton is of particular interest due to its direct response to nutrient level variability or other disturbances through changes in biomass and composition (e.g. Moustaka-Gouni 1993; Ptacnik et al. 2009).

Problems in aquatic ecosystem management include insufficient data quantity and quality, absence of systemic thinking and lack of empirical evidence for important mechanisms that presumably modulate lake patterns (Hammer et al. 2011; Vlachopoulou et al. 2014; Voulvoulis et al. 2017). Lack of standardization, inconsistencies in taxonomy, uncertainties in the characterization of eutrophication status based on phytoplankton taxa, poorly defined reference conditions, heterogeneity in data sets and lake types, weak links of national types to IC types which in turn pose comparability problems of ecological status across Europe (e.g. Nixon et al. 2012; Katsiapi et al. 2016; Søndergaard et al. 2016) remain a challenge eighteen years after WFD introduction.

Departure from the original WFD theoretical underpinning and the actual implementation efforts needs to be critically evaluated across the Community. In a recent comprehensive study, Voulvoulis et al. (2017) reviewed the problems with interpretation and the implementation of the Directive, indicating a profound misunderstanding even of its core principles. Moss (2008) argued, “A *rich resource of genuine ecological expertise in the universities and non-governmental organizations has been avoided in favour of consultancy contracts controlled to deliver a politically expedient product*”. According to Nixon’s et al. (2012) overview of main and supporting competent authorities in the MSs only Denmark, Sweden and Portugal refer to universities as supporting competent authorities. Although knowledge and experience of experts is a valuable source of information in ecological assessment of lakes, there has been little formal recognition that expert knowledge can shed light on the actual implications of measured environmental and/or biological variables (Peeters et al. 2008). Administration and water management authorities depend on reliable measurements and robust assessment methods, whereas the lack of a European-wide harmonization for comparability undermines on-going environmental protection efforts and aspirations to satisfy public demands (Arle et al. 2016).

In this article, we aim to analyse knowledge gaps, inconsistencies and limitations of WFD implementation based on lessons learned from Greece. Our examples are drawn from Greek/transboundary lakes and reservoirs in Med GIG that experience eutrophication pressure, which still represents an important threat to the integrity of freshwater ecosystems and one of the main reasons that 44% of European lakes fail to meet the “good” ecological status (EEA-ETC 2012). We revisit results from WFD implementation in Greek lakes and reservoirs in Med GIG (Greece, Spain, Portugal and Cyprus) with an intention to translate them back to ecological understanding for the protection of European lakes and to ideally strengthen the science-policy dialogue. The main pillars of our attempt to address the “knowing-doing” gap are as follows:

i. enhance ecological validity and comparability of lake typology and classification methods across Europe;
ii. question the justification of reference sites and phytoplankton reference conditions and revisit the characterization of phytoplankton components in Greek/transboundary lakes and reservoirs classification methods in Med GIG (Greece, Spain, Portugal and Cyprus); and
iii. protect Greek/transboundary Balkan lakes from misclassification and especially Lake Megali Prespa, one of the oldest lakes in Europe

## METHODS

For this article, we obtained information from several sources:

i. literature search of scientific publications
ii. a search of Joint Research Center technical reports (https://circabc.europa.eu/faces/jsp/extension/wai/navigation/container.jsp)
iii. Greek lake datasets, the original physical-chemical and phytoplankton data (period: 2012-2015) from the National Water Monitoring Network (NWMN) of the Greek Special Secretariat for Waters of the Ministry of Environment and Energy
iv. Greek Report on the application of phytoplankton index NMASRP for reservoirs in Greece submitted in June 2016 and approved in October 2016 by ECOSTAT (https://circabc.europa.eu/webdav/CircaBC/env/wfd/Library/working_groups/ecological_status/05%20%20Intercalibration%20of%20Ecological%20Status/Intercalibration%20of%20new%20or%20revised%20methods/Lakes/Phytoplankton/GR_PHP_phytoplankton%20reservoirs%20report%20to%20IC%20RP%20(1).pdf)
v. Greek Report on the development of the national method for the assessment of ecological status of natural lakes in Greece using the biological quality element “phytoplankton” revised and approved in January 2017 by ECOSTAT (https://circabc.europa.eu/webdav/CircaBC/env/wfd/Library/working_groups/ecological_status/05%20%20Intercalibration%20of%20Ecological%20Status/Intercalibration%20of%20new%20or%20revised%20methods/Lakes/Phytoplankton/GR_phyto_natural%20lakes_jan%202017.pdf)
vi. reports and reviews on WFD platform CIRCABC (https://circabc.europa.eu)
vii. direct contact with ECOSTAT members, WFD Intercalibration (IC) coordinator and Greek Special Secretariat for Waters of the Ministry of Environment and Energy
viii. Unpublished data of the authors

## LAKE TYPOLOGY

### General

In his classic lake typology, Thienemann (1925, 1927) distinguished oligotrophic and eutrophic lakes by their morphometric and hydrographic properties. Small and shallow lakes with broad littoral banks and a high perimeter: area ratios are more often conducive to eutrophic conditions, while large and deep lakes with narrow littoral banks and a low perimeter: area ratio are often characterized by oligotrophic conditions. Meanwhile, anthropogenic eutrophication and its regional variability disconnected this equivalence between morphometry and trophic state. Even a lake as big as Lake Erie (25,744 km^2^) became impacted by harmful cyanobacterial blooms (Brooks et al. 2016). On the other hand, small high elevation mountain lakes with catchment areas containing little soil and mainly bare rock can be oligotrophic (Lampert and Sommer 2007). Still it remains that small low-or midland lakes can be eutrophic even under pristine conditions and, therefore, a reference state other than large lakes is needed. During the period of intensive eutrophication studies in the 1970s, a suite of different morphological and hydrographic properties were related to the sensitivity of lakes to eutrophication, such as lake volume, lake area, lake mean depth, drainage basin area, retention time, and diverse ratios between these fundamental properties (Dillon and Rigler 1975; Vollenweider and Kerekes 1982). Nutrient loading (per unit are or volume) was defined as a key process linking nutrient export from the watershed to the nutrient inventory of a lake. Smaller lakes receive higher areal or volumetric loading at the same level of nutrient export. In addition to external loading, internal loading of the epilimnion from the sediment will be stronger in non-stratified or stratified lakes with high ratio of the area of the epilimnion sediment to epilimnion volume (Fee 1979). Whether a lake stratifies depends on both the maximum lake depth and the surface area, whereas thermocline depth (epilimnion volume) depends primarily on lake surface area (Gorham and Boyce 1989). The importance of depth is also reflected in the morphoedaphic index (MEI) used to predict fish yield of lakes (Ryder 1965).

Lakes morphometry is not only important for the response of in-lake nutrient inventories to loading, it also has direct implications for several ecosystem functional and structural properties, either as a stand-alone factor or in conjunction with lake productivity, such as the extent of physiological P-limitation (Guilford et al. 1994) because of increased mixing depth and, therefore, a higher vertical upward transport form nutrients and shift towards light imitation with increasing wind fetch. Guilford et al. (1994) analysed several physiological indicators of P-limitation of phytoplankton (seston C:P and N:P-ratios, C:chlorophyll ratio, alkaline phosphatase activity) in 9 Canadian, P-deficient lakes from ca. 0.3 to 82,000 km^2^ (all stratified with water renewal time >5 yr) and found a decrease of P-limitation with increasing area, although the response began to flatten at intermediate areas (around 10 -20 km^2^), depending on the P indicator chosen. A weaker physiological limitation by a nutrient means that less biomass is built up per unit nutrient (Droop 1973). Along the same line of thinking, Staehr et al. (2012) studied several measures of whole-ecosystem metabolism in 25 meso-to hypertrophic Danish lakes with surface areas from 0.001 to 17 km^2^ and mean depths from 0.5 to 13.5 m and found that gross primary productivity (GPP) and ecosystem respiration (R) showed a negative response to depth while net ecosystem productivity (NEP = GPP-R) showed a positive response to area and a negative response to percentage forest cover along the shoreline.

The positive relationship of species richness and ecosystem size, as shown by the species-area curve, is one of the cornerstones of ecology. For lakes, such a positive relationship between lake size and species richness has been reported for fish (Griffith 1997), macrophytes (Rørslett 1991), and zooplankton (Dodson et al. 1992). Surface area appeared important in favoring the dominance of particular species in an analysis of phytoplankton data from about 1500 lakes in twenty European countries (Maileht et al. 2013). Dodson et al. (2000) reported a significant interplay between productivity and lake area. In particular, an optimum of phytoplankton species richness was found at intermediate productivity for lake areas <100 km^2^, while a transition to a slightly U-shaped pattern with an overall declining trend was registered for lake areas >100 km^2^. In a similar manner, Smith et al. (2005) analyzed data from 142 different natural ponds, lakes, and oceans and 239 experimental ecosystems that revealed a strong phytoplankton species-area relationship. Similar to taxon diversity within functional groups, lake size also influences the number of trophic levels. Using stable isotopes as a trophic level proxy, Vander Zanden et al. (1999) found an increase of the trophic level of the top predator (lake trout) with fish diversity and lake area, except for the largest lakes (Lake Ontario, Lake Superior) in the data set. They argued in favor of “productive space” (area x productivity) as a more sensible predictor for food chain length. By contrast, Post et al. (2000) showed a linear increase of food chain length with lake volume, casting doubt on the ability of productivity and productive space to be used as predictors. The aim of lake typology is to produce a simple, ecologically-relevant classification scheme that effectively balances the degree of specificity (or detail) with ease of implementation. Increased granularity, i.e. more descriptors of typology and finer scale thresholds can conceivably provide a better representation of all the influential factors, but if the types become too narrowly defined, the number of lakes per type diminishes, reducing the likelihood of finding a reference (i.e. pristine) lake in each type. Moreover, “natural” limits (gaps in frequency distribution, breakpoints in response curves) should be preferred over arbitrary ones, but are often difficult to find, because of continuous responses or because different ecological response variables suggest different limits (see citations above). The only obvious limit is the one between polymictic and summer stratified lakes, which obviously depends on depth. However, the depth limit between both types depends also on local climate, lake length/wind fetch and water transparency (Read et al. 2014; Kirillin and Shatwell 2016). But even in the case of stratification type ambiguity arises when a lake with complex morphology consists of stratifying and polymictic basins. Moreover, the traditional mid-lake or deepest point-sampling misses water quality differences between the different lake basins, which in extreme cases should be dealt as separate water bodies while shoreline features can comprise a hydraulic storage zone that enhances harmful algal persistence (e.g. Grover et al. 2010).

The difficulties in choosing appropriate depth and area boundaries could be circumvented by using two alternative descriptors, which capture the two dominant biogeochemical issue related to morphometry, sediment -surface water interactions and land/shore based influences on the lake: the percent share of epilimnion area water in contact with the sediment (by definition 100% in non-stratifying lakes) and the lake volume: watershed area ratio. Values for type boundaries will have to be decided in the discussion and decision processes during future modifications of the WFD whereas systems thinking is a pre-requisite to effective WFD implementation (Vlachopoulou et al. 2014; Voulvoulis et al. 2017).

We emphasize, that our typological considerations are tailored to fresh-water and clear-water lakes, while oligo-haline, brown-water and silted lakes need special treatment, because their specific physico-chemical properties affect biological elements. Accordingly, WFD (ANNEX II and V) requests transparency and salinity be included in the physico-chemical properties that shape the biological response, while the mean substratum composition is considered as an optional factor of typology system B (European Commission 2000).

### Lake typology in GIGs and Greece

Greece belongs to the Med GIG, started its WFD implementation work in 2004 and the first results for lakes were published in 2008 (phytoplankton metrics and their boundaries). The next phase results for lakes were published in 2014 (phytoplankton methods and their boundaries) (de Hoyos et al. 2014). Greece participated in the collection of intercalibration data set only with one reservoir (Tavropos) assigned preliminarily by the Med GIG to the Mediterranean type LM5/7 (deep, large, siliceous with low alkalinity). Phytoplankton metrics and the reference conditions (as site-specific not type-specific) for Tavropos Reservoir were provided to the Med GIG by the first author of the present work (see Phillips et al. 2013; Pahissa et al. 2015). The next step followed throughout the WFD implementation in Greece was the 2012-2015 NWMN. In the following, the proposed national typology of Greek lakes resulting from this monitoring effort will be revisited.

The flexibility allowed by the WFD for MSs to develop national methods and typologies has resulted in a wide variety of classification schemes (ca. 673 lake types; Nixon et al. 2012; Poikane et al. 2014). After the IC exercise, only one third of the national lake types were linked to common IC types (Solheim et al. 2011). Even with the reduced classification scheme significant variation exists in selecting descriptors in different GIGs (Pardo et al. 2011). The common biogeographical types across Europe were insufficiently standardized to determine consistent, type-specific reference conditions (Poikane et al. 2014). A characteristic example is the omission of surface area as a lake type descriptor by IC exercise (e.g. Pahissa et al. 2015; Table 2) although a high number (18) of MSs selected this as an important underlying factor (Solheim et al. 2011). A lack of explicit consideration of surface area boundaries for large lakes was indirectly addressed by the adoption of a threshold value of 0.5 km^2^ to delineate small lakes (e.g. Wolfram et al. 2014). This is not explained by known lake functional and structural properties (i.e. Guilford et al. 1994) while a high number of lakes (1150) within the EU territory have surface areas larger than 10 km^2^ (Tsavdaridou pers.com.).

Correspondence between the IC typology of reservoirs and national typologies can be problematic (de Hoyos et al. 2014). In the Mediterranean GIG, all the reservoirs considered had a mean depth >15 m, a surface area >0.5 and < 50 km^2^, a catchment area <20,000 km^2^, and were intercalibrated based on different alkalinity/geology and climatic factors (Pahissa et al. 2015). In the same context, Greece recently applied the New Mediterranean Assessment System for Reservoirs Phytoplankton (NMASRP), a multiparametric index composed of four parameters (chlorophyll *a*, total biovolume, IGA Index Des Grups Algals, biovolume of cyanobacteria) (de Hoyos et al. 2014). Twenty reservoirs in Greece were assigned to LM5/7 (deep, large, siliceous with low alkalinity) and LM8 (deep, large, calcareous with high alkalinity) common types of the Mediterranean GIG (Table S1). However, NWMN data (Table S1) suggest that the reservoirs assigned to LM5/7 are characterized by high alkalinity values characteristic of LM8. It is also worth noting that large and deep Greek reservoirs for which the Mediterranean GIG was applied had a wide range of surface areas and mean depth, varying from 0.47 km^2^ and 10.5 m (Techniti Limni Feneou) to 66.6 km^2^ and 46.7 m (Techniti Limni Kremaston) (Table S1). According to the Med GIG scheme (surface are <50 km^2^), Techniti Limni Polyphytou and Kremaston should not be assigned either to LM5/7 or LM8. In addition, retention time or flushing rate should have been considered as a fundamental type descriptor, because water bodies with faster water renewal can assimilate a larger phosphorus load with no adverse eutrophication responses compared with slower-flushing water bodies (Lampert and Sommer 2007; Pardo et al. 2011). Flow in riverine reservoirs can be strong enough to wash out phytoplankton populations, including harmful algae (Grover et al. 2010).

Greek national typology splits natural lakes only by mean depth using 9 m as a threshold for deep stratified lakes. This threshold, not linked to any IC type of the GIGs (e.g. de Hoyos et al. 2014; Wolfram et al. 2014), was a failed attempt to distinguish between stratification regimes (polymictic and warm monomictic). In fact, the depth separating polymictic from monomictic / dimictic lakes also depends on altitude, lake length, wind shelter, water transparency and local climate, as shown by recent, physics-based analyses by Read et al. (2014) and Kirillin and Shatwell (2016). The failure of the 9 m threshold can be seen by comparing two similar shallow and similar sized transboundary lakes of different altitudes in the Mediterranean region, Lake Mikri Prespa (4.7 m deep, 854 m ASL, surface area 48 km^2^) is polymictic (Tryfon et al. 1994), whereas Lake Doirani (4.3 m deep, 146 m ASL, surface area 30 km^2^) tends to stratify during the summer months (Temponeras et al. 2000). For this reason, we suggest using actual stratification regimes of shallow lakes instead of a fixed threshold.

Lake assignment into national types. When designing national typology, the mandate of MSs is to focus on the overall purpose of the Directive outlined in Article 1, i.e. establish a framework for the protection of inland and other waters, prevent further deterioration, and protect and enhance the ecosystem status. Typology is the tool to assist the Article 1 goal by facilitating the impartial relative assessment of comparable ecosystems. Using defensible values for water quality descriptors, lake typology should ensure that appropriate phytoplankton reference sites with type-specific baseline conditions are selected in order to offer a reliable ecological status classification of non-reference lakes with similar basic characteristics (e.g. surface area, altitude, salinity, turbidity). Here, we will use the example of Greece to illustrate our doubts about the scientific validity of the practices followed. Lakes Kourna and Paralimni (Table 1) have been selected as phytoplankton reference sites for deep and shallow lakes, respectively, while lake size and any other basic features like altitude, salinity, climate factors, inherent detritus water content have not been considered as typology descriptors. Furthermore, descriptors that capture systems biogeochemical issues related to morphometry such as sediment -surface water interactions and land/shore based influences on the lake have also not considered. As shown in Table 1, the ecological-grouping of lakes is highly questionable as it clusters together systems with significant differences in their basic limnological and climatic characteristics. Based on evidence provided in Table 1, Figures 1-3 and Tables S2, S3 Kourna and Paralimni cannot serve as reference sites for the large lakes of Greece and transboundary Balkan lakes. As will be discussed in the following section, the biological justification of the selected reference sites is also profoundly problematic.

**Figure 1.**
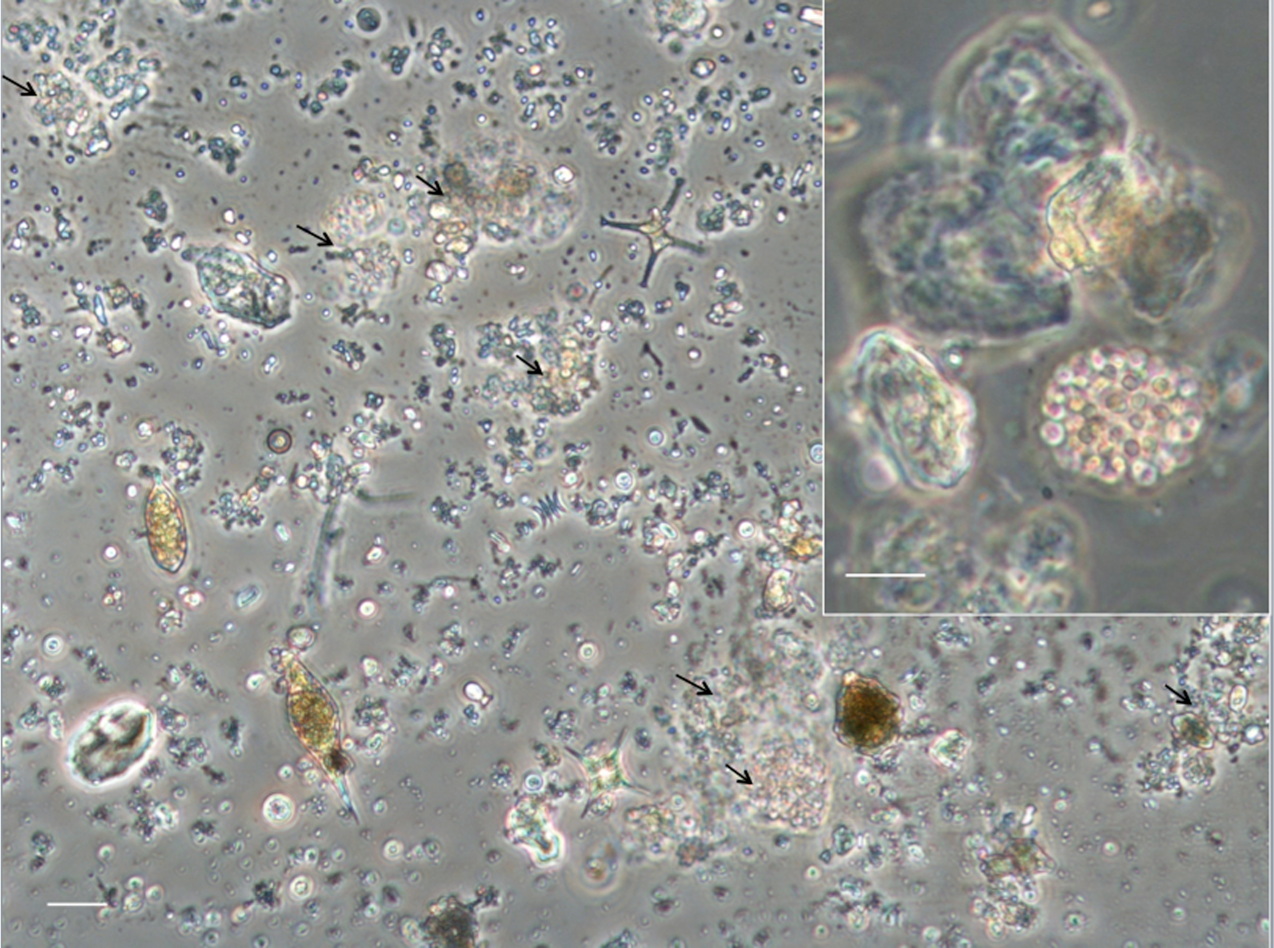
Micrograph of a water sample of Lake Paralimni in July 2017: phytoplankton individuals, aggregates of colonial small-celled cyanobacteria and non-living suspended particles. Black arrows show aggregates. Scale bar is 20 µm. Insert: An aggregate in magnification. Scale bar is 10 µm.

**Figure 2.**
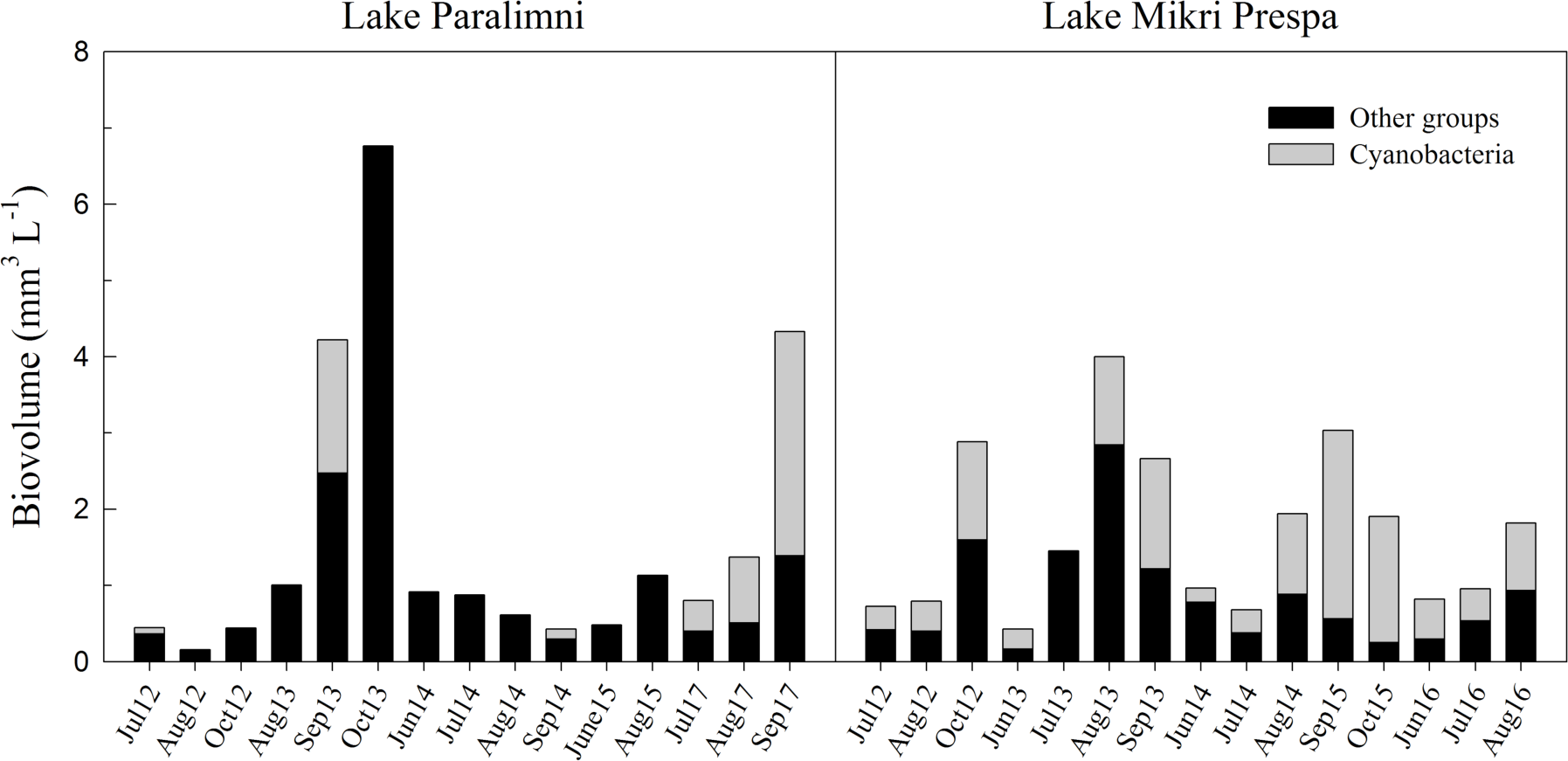
Temporal phytoplankton and cyanobacteria biovolume variations in Greek shallow lakes, Paralimni (reference site) and Mikri Prespa (moderate ecological status).

**Figure 3.**
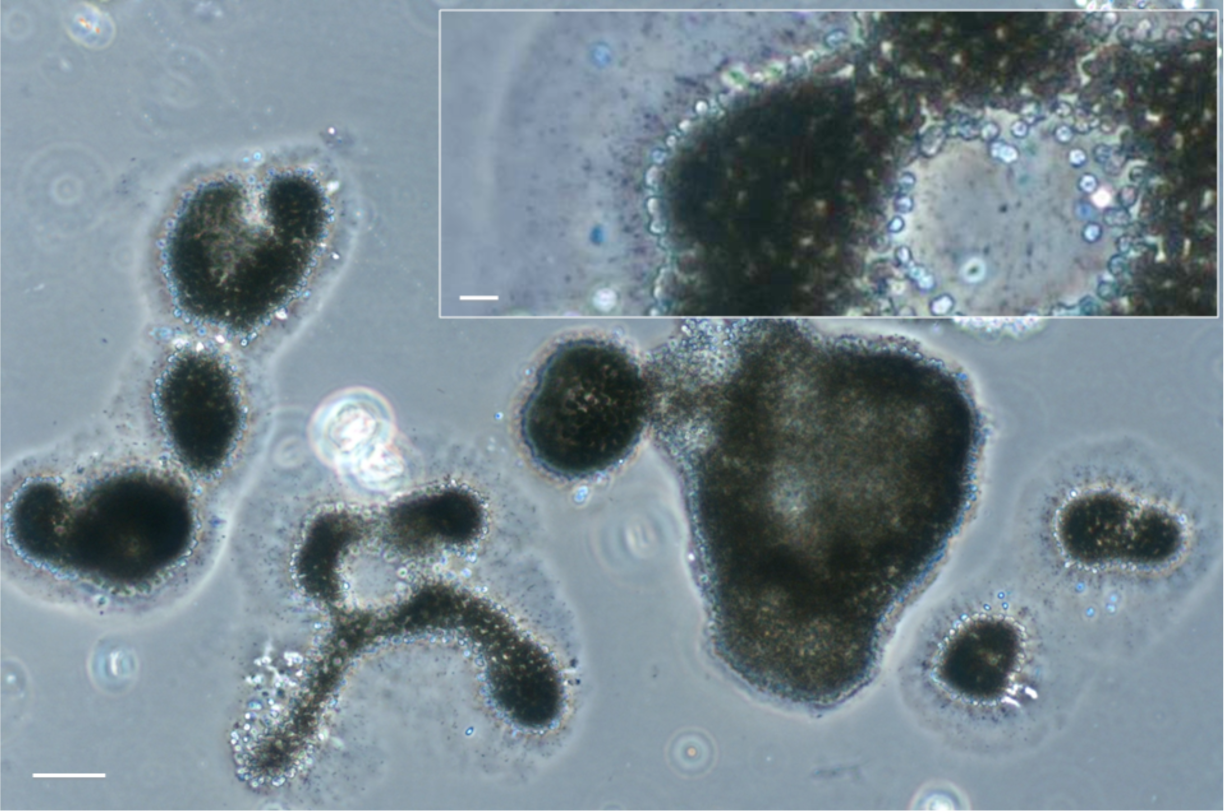
Micrograph of a water bloom sample of Lake Paralimni in September 2017. *Microcystis* colonies attacked by bacteria. Scale bar is 50 µm. Insert: A magnification of attacked *Microcystis* colony. Scale bar is 10 µm.

**Table 1.**
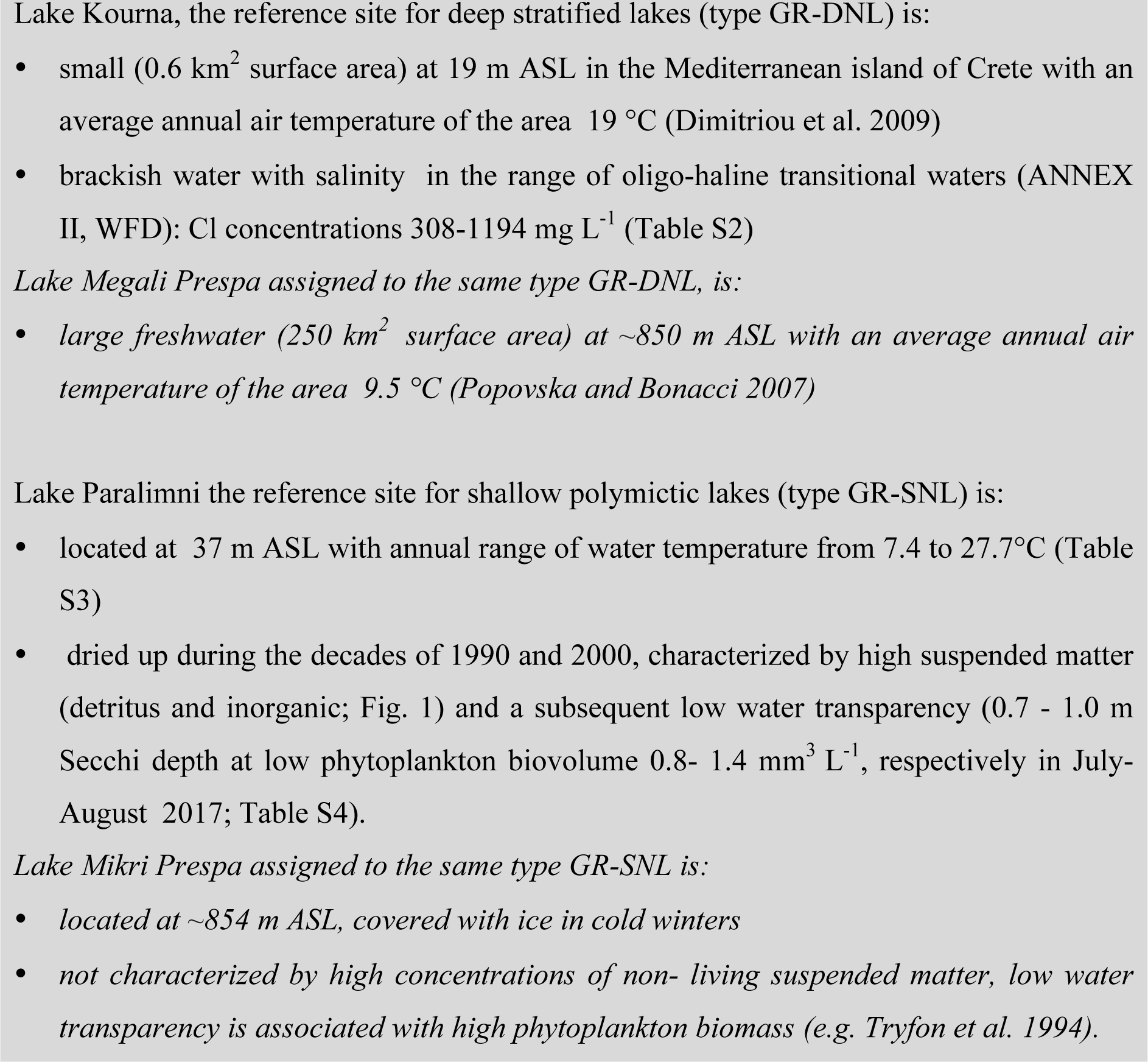
Evidence for ecologically invalid national lake typology.

## UNDERSTANDING THE ECOLOGY BEHIND PHYTOPLANKTON METRICS AND INDICES

### Ecological gaps associated with phytoplankton and cyanobacteria biovolume

The selection of two highly questionable reference sites led to the paradoxical situation, in which the reference value for total phytoplankton biovolume in deep lakes became higher than that for shallow lakes (1.29 mm^3^ L^-1^ in deep lakes; median of 7 values from the warm period based on two years of data from Lake Kourna) as opposed to 0.74 mm^3^ L^-1^ in shallow lakes (median of 12 values from the warm period based on four years of data from Lake Paralimni) (Tables S4, S5). This delineation of the baseline conditions contradicts widespread limnological experience, as well as the reference definitions by other MSs across the GIGs (Table S6). For example the reference values for Austria are 0.3 mm^3^ L^-1^ for deep L-AL3 lakes (annual means) and 0.7 mm^3^ L^-1^ for shallower L-AL4 lakes (Wolfram et al. 2009). While the higher values set for Greek lakes may partly stem from the warm period values and overly ambitious Austrian targets, the inversion between Greek shallow and deep lakes is void of any ecological plausibility.

Phytoplankton has been sampled during 2012-2015, four consecutive years for most lakes, but in some cases only during three years (e.g. Prespa lakes and Lake Kourna). For the reference Lake Kourna, one of the three sampling years (2014) has been excluded from the analysis (Table S5), without obvious scientific reason. Noteworthy is that the excluded phytoplankton biovolume values were higher (range from 1.28 to 6.54 mm^3^ L^-1^) than those from the two remaining years (Table S5) included in the analysis, indicative of significant interannual variability in phytoplankton biomass, as is usually the case for other lakes (e.g. Moustaka-Gouni 1993; Roelke et al. 2004). Inclusion of the year 2014 would have led to an even higher reference phytoplankton biovolume value (2.0 mm^3^ L^-1^instead of 1.29 mm^3^ L^-1^, almost three times the reference value of shallow Greek lakes), which reinforces our point that Lake Kourna is absolutely unsuitable as a reference site (Table 2). The reference phytoplankton biovolume levels for large, deep Greek reservoirs, assigned into two IC types LM5/7 and LM8 (Table S1) in application of the NMASRP index (Greece, Cyprus and Portugal), are 1.2 and 0.9 mm^3^ L^-1^ (Table S6), respectively (de Hoyos et al. 2014). Given that the lower retention time in reservoirs compared to lakes should lead to a weaker biovolume response to nutrients (Lampert and Sommer 2007), it can be easily inferred that the practices followed in Greece have resulted in counter intuitively higher reference values relative to the one established for shallow Greek lakes (0.74 mm^3^ L^-1^). Compared to other Mediterranean countries using the same metrics/indices (Table S6), the reference phytoplankton biovolume of 1.2 mm^3^ L^-1^ for Tavropos Reservoir of LM5/7 IC type used by Greece, Cyprus and Portugal is much higher than the 0.36 mm^3^ L^-1^ reference phytoplankton biovolume used for the same IC type LM5/7 by Spanish MS in the MASRP index (de Hoyos et al. 2014). Interestingly, the 0.36 mm^3^ L^-1^ value was originally derived from the Tavropos Reservoir data and provided by the first author of this work to the IC data set (see Maileht et al. 2013; Pahissa et al. 2015), but Greece applied the phytoplankton index NMASRP (used in Cyprus and Portugal) and not the MASRP (used in Spain) and the adopted reference-value ended up being four times higher for the same reservoir. Insufficient clarification of the methods chosen by authorities and insufficient evaluation of alternative indices have also been reported from other MSs (e.g. Stoyneva et al. 2014). Comparing the reference phytoplankton biovolume levels for deep reservoirs of the three Mediterranean MSs (Greece, Cyprus and Portugal) with the reference values of deep/stratified lakes in other GIGs (Table S6) shows higher biovolume values in reservoirs, although their expected lower retention time should lead to lower biovolume because of faster washout.

**Table 2.**
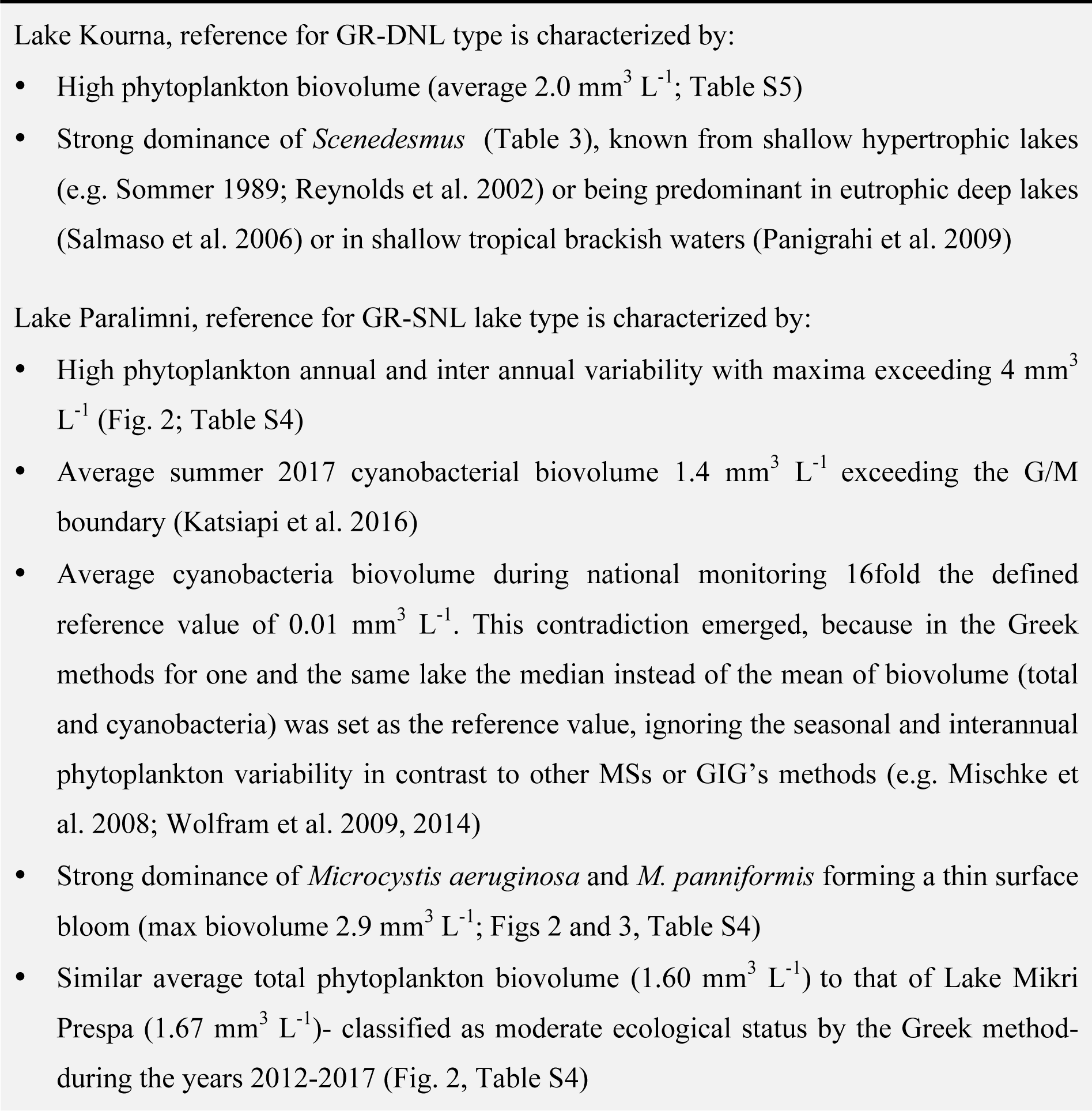
Evidence for unjustified reference phytoplankton sites/conditions.

According to WFD Annex V (European Commission 2000), frequency and intensity of planktonic blooms are important facets of phytoplankton dynamics. Cyanobacteria biovolume or their percentage contribution to total phytoplankton biovolume has been a recommended taxonomic or bloom metric for MSs of GIG’s (e.g. de Hoyos et al. 2014; Solheim et al. 2014) because of the well-known relationships between cyanobacterial blooms and eutrophication and the adverse effects of cyanobacterial blooms on water quality and ecosystem services (e.g. Brooks et al. 2016). However, not all cyanobacteria are included in the Greek classification method for lakes and reservoirs as well as in the Med GIG assessment methods for reservoirs (de Hoyos et al. 2014) in contrast to the methods of MSs of other GIGs, except the method of UK in the Northern lakes GIG (Solheim et al. 2014). In particular, all genera of the Chroococcales except for *Microcystis* and *Woronichinia* spp. were excluded in the Mediterranean GIG methods (de Hoyos et al. 2014). *Microcystis* species were deemed an essential surrogate of eutrophication severity due to their dominance and bloom formation in eutrophic water bodies. However, significant ecological information (early warning signals of on-going degradation) can be lost by excluding other chroococcal cyanobacteria species forming blooms in mesotrophic or slightly eutrophic lakes. For example, recurring blooms of small-sized chroococcal cyanobacteria dominated by *Aphanocapsa*, typically found in eutrophic waters, have been reported in Prespa lakes (Tryfon et al. 1994; Reynolds et al. 2002; Katsiapi et al. 2012). Further, *Woronichinia*, known from eutrophic and mesotrophic lakes in central -north Europe (e.g. Wilk-WoŹniak et al. 2003), is abundant together with *Snowella* in oligotrophic and mesotrophic Finish lakes (Rajaniemi-Wacklin et al. 2005). In Greece, *Woronichinia* is rare in lakes over the entire trophic spectrum, while the sister-genus *Snowella* is among the dominant genera in eutrophic lakes (Moustaka-Gouni 1993; Tryfon et al. 1994; Moustaka-Gouni et al. 2010). In addition, some *Aphanocapsa* and *Microcystis* species, commonly observed in Greek lakes, are mere synonyms (e.g. the previously named *Microcystis incerta*, and now known as *Aphanocapsa incerta*) and/or have substantial overlap in cell/colony dimensions and morphology, while other excluded species of chroococcal genera (*Cyanodictyon, Aphanothece, Radiocystis, Merismopedia, Sphaerocavum*) are abundant in mesotrophic and eutrophic lakes (Hindak and Moustaka 1988; Tryfon et al. 1994; Reynolds et al. 2002). Therefore, the unjustified exclusion of several important chroococcal species from the assessment of cyanobacteria biovolume used to characterize phytoplankton blooms in Greece, Cyprus and Portugal can potentially undermine our ability to detect the very critical “moderate” ecological conditions.

### Problems associated with phytoplankton indices

For Greek reservoirs as for other Mediterranean MSs (Spain, Portugal, Cyprus), the index IGA (Table S7), originally developed for Catalonian reservoirs (Catalan and Ventura 2003) was adopted as a phytoplankton compositional index. This seems to be a plausible choice in principle due to the similarity of the prevailing conditions in the Mediterranean region (Moustaka-Gouni et al. 2014). However, there are problems with the inclusion of at least four indicator phytoplankton groups (colonial chrysophytes, colonial diatoms, non-colonial diatoms, dinoflagellates) and with the weighting factor of one group (colonial Volvocales). Colonial chrysophytes are not indicative of eutrophication, e.g. *Dinobryon* is also dominant in most oligotrophic Greek reservoirs (Moustaka-Gouni and Nikolaidis 1992). The association of colonial diatoms with eutrophication might be true for *Aulacoseira*, while genera of the Fragilariaceae family (*Asterionella, Fragilaria, Tabellaria*) can be abundant throughout the spectrum from oligotrophic to eutrophic lakes and reservoirs, e.g. *Asterionella gracillima* is dominant in the oligotrophic reservoir Tavropos (Moustaka-Gouni and Nikolaidis 1992), a reference Greek site. Colonial Volvocales are correctly assigned to the numerator but the high weighting factor (3, as compared to 4 for the Cyanobacteria) seems unjustified, because none of their blooms qualifies as HAB (harmful algal bloom). Including dinoflagellates into the denominator mischaracterizes the occurrence of blooms of *Ceratium* and *Peridinium* in eutrophic reservoirs (e.g. Hart and Wragg 2009; Kang et al. 2008), the *Ceratium* blooms in eutrophic lakes (e.g. Temponeras et al. 2000; Reynolds et al. 2002), the *Peridinium gatunense* blooms in meso-eutrophic Lake Kinneret (Zohary et al., 1998), and the *Peridinium*-*Woronichinia* association in stratified mesotrophic lakes (Reynolds et al. 2002). The position of non-colonial diatoms in the denominator is also problematic. It is justified for some solitary *Synedra* spp. but not all (e.g. *Synedra acus* known from eutrophic shallow lakes and rivers), while the presence of *Cyclotella* spp. and *Stephanodiscus* spp. along the oligo-to eutrophic continuum is also species-specific (Sommer 1985; Sommer et al. 1993; Reynolds et al. 2002). Instead of the IGA index, it is evident that species-specific indicators of composition metrics may be more appropriate (e.g. Marchetto et al. 2008 in Italy), taking into consideration the tolerance to flushing as a key phytoplankton trait for reservoir community assembly and the reservoir phytoplankton as an intermediate between lake plankton and potamoplankton. On the other hand, the high weighting factor for cyanobacteria resulting in high IGA values whenever a reservoir experiences a cyanobacterial bloom does not add value to the simply assessed bloom by calculating cyanobacteria biovolume. There is also a contradiction between the two phytoplankton compositional descriptors (community description and IGA index) used in the Mediterranean GIG methods regarding the taxa used as eutrophication indicators in LM5/7 reservoirs (see de Hoyos et al. 2014). For instance according to the community description, reservoirs at Maximum Ecological Potential in the Mediterranean are dominated by the colonial chrysophyte *Dinobryon*, colonial chlorococcales (*Sphaerocystis* and *Coenochloris*) and the colonial diatom *Asterionella*, which are all placed in the numerator together with cyanobacteria as indicators of eutrophication.

The Nygaard index, modified by Ott and Laugaste (1996) for small Estonian lakes, was selected for the ecological classification of large Greek lakes (Table S7). Latvia (Central Baltic GIG) also used the adapted Estonian lake phytoplankton method including this modified Nygaard index (Phillips et al. 2015). In this index, the centric diatoms are placed in the numerator together with cyanobacteria as indicators of eutrophication while the presence of the most common centrales *Cyclotella* and *Stephanodiscus* along the oligo-to eutrophic continuum is species-specific (e.g. Reynolds et al. 2002). Because the diatoms *Cyclotella commensis* and *C. bodanica* from Centrales were restricted to high and good quality reservoirs, the selected index was further modified by the Greek authorities to exclude Centrales as an indicator of eutrophication according to Katsiapi et al. (2016). However, according to this modification of the Nygaard index (Table S7), cryptophytes should also be excluded (Katsiapi et al. 2016) since they are not restricted to eutrophic waters, while strong seasonality characterizes their dominance in oligotrophic large lakes (Sommer et al. 1993). Most importantly, there is a clear contradiction between the two phytoplankton compositional descriptors (community description and modified Nygaard index) used in the Greek method regarding the taxa used as eutrophication indicators in the same report. For instance according to the phytoplankton community description, high ecological status Greek lakes are dominated by the chlorococcales *Oocystis* and *Monoraphidium*, the cryptophytes *Cryptomonas* and *Rhodomonas* along with a small number of chrysophytes (Table 3), while chlorococcal genera can also be encountered in lakes of good ecological status. However, in the modified Nygaard index, Chlorococcales and Cryptophyta (which include the above mentioned genera) are placed in the numerator together with cyanobacteria and euglenophytes, as indicators of eutrophication. Consequently, the application of the Greek modified Nygaard index on phytoplankton communities can lead to mischaracterization of lake ecological status relative to existing empirical evidence.

**Table 3.**
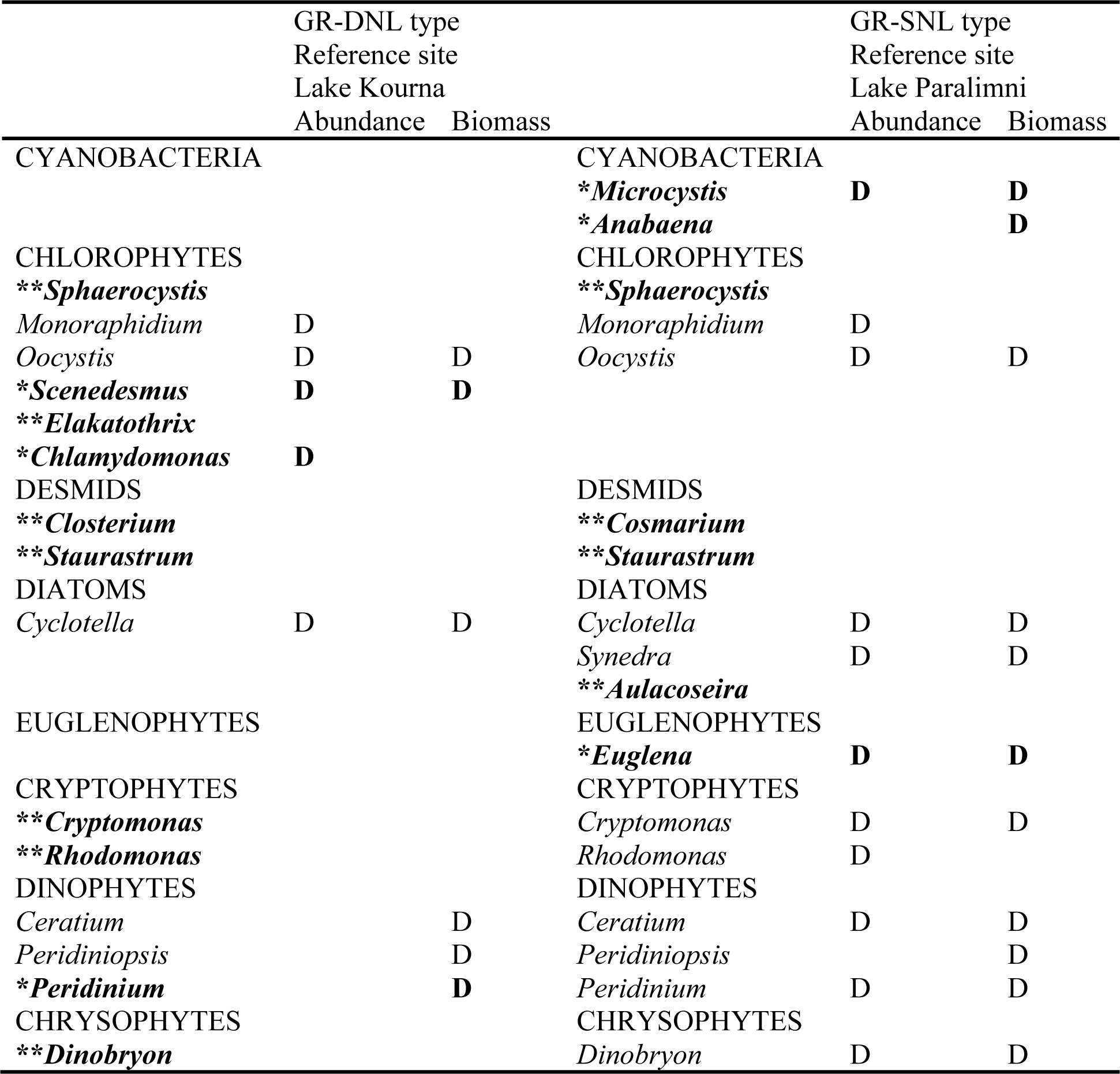
List of phytoplankton genera: a) dominant with contribution >10 % to the total abundance or biovolume in each sampling of the reference sites Lake Kourna (type GR-DNL) and Lake Paralimni (type GR-SNL) of the national monitoring data set and b) which are presented as the descriptors of the phytoplankton community of the high status in the Greek classification method on the basis of the same national monitoring data set. In Bold: the wrong genera. Bold with one asterisk: genera of (a) not included in (b). Bold with two asterisks: genera of (b) not included in (a). D: dominant.

Major phytoplankton genus/species level changes occurring during seasons are widespread in lake phytoplankton. Therefore, the description of phytoplankton communities for high, good and moderate status presented in reports is not a reliable status indication because of insufficient coverage of seasons. Moreover, there is a great discrepancy between the genera in described communities at high status in both deep and shallow lakes and the genera dominant in abundance or biovolume in the reference sites Lake Kourna and Lake Paralimni, all genera/data based on one and the same national monitoring (NWMN) data set (Table 3).

PhyCoI, a new phytoplankton community index suggested by Katsiapi et al. (2016) for assessing ecological water quality and tested by research data from Greek lakes and reservoirs was rejected by the Greek authorities because of a formal lack of compliance with WFD requirements. The lack of compliance according to the Greek reports was that no conversion of the 0-5 scale of the five ecological classes into the scale of 0-1 of EQR was provided and the scoring was made by expert judgment based on spatial data. While the conversion from a 0-5 to a 0-1 scale could have easily been made, Greek authorities refused expert judgment (but see Baattrup-Pedersen et al. 2017). This index is calculated from the scores of five different metrics/sub-indices including two sub-indices of biodiversity metrics, which is among the priority issues identified for enhancing WFD implementation (Reyjol et al. 2014). The Greek ecological classification method for lakes is composed of four metrics(chlorophyll *a*, total biovolume, modified Nygaard index, biovolume of cyanobacteria), that(except chlorophyll) have been critically reviewed above. The establishment of the Greek method was based on regression analyses of the developed index from 2014 and 2015 phytoplankton data against TP concentrations from 2015 alone, i.e. the same TP values were apparently regressed against the phytoplankton data from 2014 (Table S8). This double use is conceptually illegitimate and statistically invalid, because of non-independence of the data used for the predictor variable. Lakes Kourna and Paralimni, which as made abundantly clear in our paper, are not appropriate types for determining the status of large deep and shallow Greek/transboundary lakes and do not qualify as representing reference conditions. In doing so, the protection targets of Greek/transboundary Balkan lakes are unjustifiably relaxed and introduce comparability problems of ecological status across Europe, even though the cognitive basis of the WFD amply demonstrates how knowledge is supposed to support the definition, monitoring and assessment of water management actions (Steyaert and Ollivier 2007).

## THE PARADIGM OF LAKE MEGALI PRESPA, A EUROPEAN ENDAGERED MONUMENT NEEDING PROTECTION

Lake Megali Prespa is a transboundary lake shared by Greece, the Former Yugoslavian Republic of Macedonia (FYROM) and Albania. It is a monomictic lake located at ∼850 m above sea level with a surface area of ∼250km^2^ (Table 1; Katsiapi et al. 2012), maximum depth ∼50 m, and mean depth ∼20 m (Löffler et al. 1998). Lake Megali Prespa has been designated as part of the largest national park in Greece and is covered by the Ramsar Convention (http://www.ramsar.org), as it hosts a significant number of rare and endemic species (Albrecht and Wilke 2008). However, over the last few decades there has been increasing concern over its ecological status due to ongoing eutrophication and water level decline. Based on the NMWN phytoplankton biovolume data (2012-2014; Table S9) and using the boundaries for status classes and the type -specific functions for stratified lake types (IC relevant lake types AL3 and AL4) according to Mischke et al. (2008) Lake Megali Prespa is classified as lower than good ecological status (Table S9). Phytoplankton biovolume (3.85 mm^3^ L^-1^) was measured in the lake in 2008, indicative of a lower than good water quality (Katsiapi et al., 2012). The phytoplankton community index PhyCoI (Katsiapi et al. 2016) using the most recent phytoplankton data (2015-2016; Katsiapi et al. in prep.) is indicative of moderate ecological water quality. Furthermore, cyanobacteria made a high contribution (29.7 -60.0%) to the annual total phytoplankton biovolume during the years 2012 – 2014 (Table S9). In June 2016, a cyanobacterial bloom occurred in Lake Megali Prespa, dominated by the species *Anabaena/Dolichospermum lemmermanii* (Figure 4), known for its ability to produce various toxins (e.g. microcystins, anatoxins, saxitoxin) (Lepistö et al. 2005). It has been confirmed as an anatoxin producer and cause of bird kills in Danish Lakes (Onodera et al. 1997).

**Figure 4.**
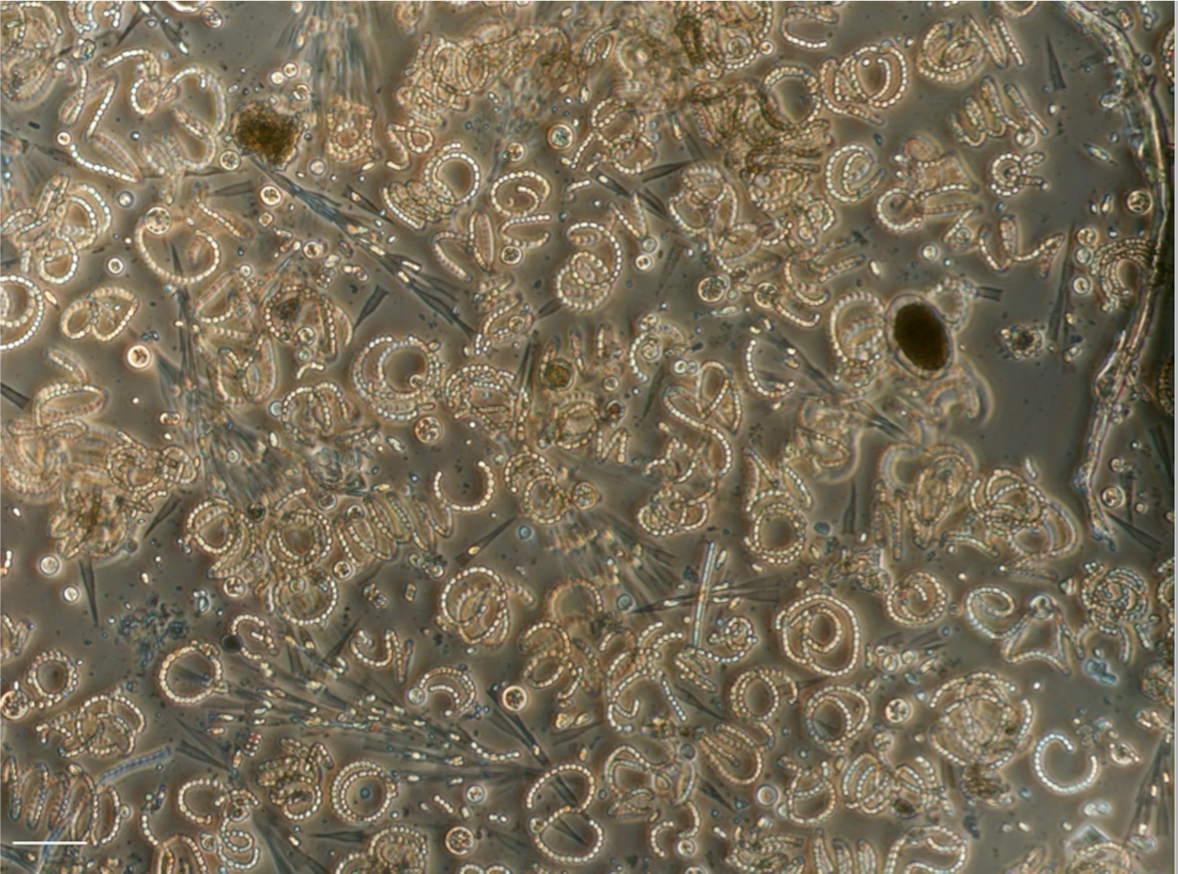
Micrograph of water bloom sample of Lake Megali Prespa in June 2016. *Anabaena lemmermanii, Dinobryon bavaricum* and *Cyclotella* species are shown. Scale bar is 50 µm.

Prespa lakes is a well-known location in Europe for bird breeding and overwintering due to its high richness and internationally important endangered species (Catsadorakis 1997), holding the largest population of Dalmatian pelicans in the Mediterranean (Catsadorakis et al. 2015). The Dalmatian pelican (*Pelecanus crispus*) is currently a species of global conservation concern, listed as “Vulnerable” in the IUCN Red List (IUCN 2014). Cyanotoxins were a plausible cause for this bird’s mass mortality episode in the Greek Karla Reservoir in 2016 (Papadimitriou et al. 2018). Therefore, such cyanobacterial blooms pose threats for the rich, unique and vulnerable bird life of the lake. In the same context of water quality degradation, Salmaso et al. (2015) identified a strong association between the shift of Lake Garda (large and deep lake south of Alps) from ultraoligotrophy-oligotrophy to oligo-mesotrophy and the development of *D. lemmermannii* surface blooms. The biovolume value of the same species in the trophogenic layer of Lake Megali Prespa in 2016 was 50 times higher than the corresponding biovolume in Lake Garda indicating higher than mesotrophic conditions. Moreover, in Lake Megali Prespa, a persistent *Anabaena* toxic bloom occurred during May -September 2010in the area of the lake close to the Greek -FYROM borders (Krstić 2012). Therefore, the recorded phytoplankton blooms in the lake during 2008-2016 are obvious symptoms of eutrophication, calling for action by the local and national water authorities. Further, functional metrics/indices of food-web, i.e. zooplankton grazing potential (although not included in the biological elements of WFD) showed that Lake Megali Prespa is undergoing eutrophication and water quality degradation (Katsiapi et al. 2012; Katsiapi et al. in prep.). The overall status of the lake that the WFD seeks to improve rather than the individual element phytoplankton outlined in Annex V of the Directive is required (see Voulvoulis et al. 2017).

All the above mentioned plankton attributes in combination with the strong lowering of mean annual water transparency in the deep and large Lake Megali Prespa, 4.4 to 2.2 m during 2012-2016 (Table S10) as opposed to 10.0 to 7.2 m during the 30s and 50s (Löffler et al. 1998), the recent summer anoxia and accumulation of phosphate phosphorus and ammonia nitrogen above sediment (Matzinger et al. 2006) are signs of degradation of high and good water quality. The ecological health and integrity of Lake Megali Prespa is further deteriorated by water losses to irrigation, which have been shown to accentuate the severity of eutrophication phenomena and may put at risk the future status of the neighboring Lake Ohrid (Matzinger et al. 2006). Yet, in the revised RBMP of Greece in 2017 based on the NWMN (http://wfdver.ypeka.gr/wp-content/uploads/2017/08/EL09_1REV_P13_Prosxedia_LAP_v02.pdf), Lake Megali Prespa has been classified in good ecological status compared to the moderate ecological status classification based on expert judgment and reported in the first RBMP (http://wfdver.ypeka.gr/wp-content/uploads/2017/04/files/GR09/GR09_P26d_Perilipsi_Prespes_EN.pdf), although no evidence of water quality improvement has been reported during the 2014-2017 period. This may demonstrate how invalid reference site (Lake Kourna in Crete) and reference conditions, choice of ecologically inappropriate indices (as made clear in our paper) and lack of expert judgment can lead to misclassification of Lake Megali Prespa ecological status.

Misclassification may lead to wrong management decisions (Søndergaard et al. 2016), i.e. to omission or at least delay of necessary and sufficient restoration measures to prevent further deterioration, protect Greek lakes and especially Lake Megali Prespa, the ancient sister of Lake Ohrid.

## SYNTHESIS AND RECOMMENDATIONS

Our analysis is meant to offer a constructive critique, sensu Toomey (2016), driven by the motivation to help legal authorities to understand better how scientific shortcomings in lake typology, selection of reference sites and inappropriate biological indices may bias the outcome of lake ecological classification. The ecological misclassification of Greek/transboundary lakes and reservoirs based on phytoplankton metrics during the current WFD implementation is seen as just one example, helping to identify similar analogies elsewhere and to overcome comparability difficulties of ecological status assessment across Europe. Deficiencies are seen in all steps needed to establish a valid assessment of lake and reservoir ecological status:

1. The lake and reservoir typology of Greek/transboundary lakes and reservoirs has failed to take essential components such as surface area, salinity, content of non-living matter and retention time into account. In addition, links of national types to common intercalibration types are missing (depth) or are unclear (alkalinity).
2. The choice of reference lakes has been arbitrary, which consequently are poor representatives of their “type”. Besides being minimally impacted, reference lakes should also represent typical properties of their type in terms of surface area, altitude, salinity, suspended non-living matter, etc. rather than being outliers.
3. The phytoplankton indices originally used in other Mediterranean and Central Baltic GIG MSs partly contain erroneous assignments of phytoplankton groups to eutrophic vs. oligotrophic waters when applied to Greek lakes and reservoirs. The reference values of phytoplankton metric biovolume in Greek deep lakes and deep Mediterranean reservoirs in Greece, Cyprus and Portugal are higher compared to the reference values of the same reservoir types in Spain, of the Greek shallow lakes and of the deep and shallow lake types (most relevant) in other MSs of the GIGs. Exclusion of most chroococcal species from the assessment of cyanobacterial metric in Greek lakes and most Mediterranean reservoirs in contrast to other GIGs can potentially undermine the ability to detect on-going degradation in water quality.
4. As a consequence of the shortcomings listed above, there is limited comparability of the ecological status between Greek/transbounbdary lakes with other European lakes and the bar of quality standards has been too low in order to provide an efficient target for the protection and restoration of critically important lakes such as the ancient European Lake Megali Prespa.

We seek to overcome the deficiencies of the current practice in order to protect and restore Greek/transboundary lakes from environmental degradation, in particular eutrophication. Since the Commission plans to review the practices followed during WFD implementation, our paper intends to shed light on methodological weaknesses and knowledge gaps that undermine the ecological foundation of WFD and the efficacy of the associated remedial measures in restoring eutrophic lakes. Our recommendations are:

1. Retain depth and add surface area size as main pillars of lake/reservoir typology in order to improve comparability and ecological relevance across Europe. Add retention time as a fundamental type descriptor in reservoirs.
2. Replace fixed limits between “shallow” and “deep” lakes by the boundary between polymictic and stratifying lakes. In addition, stratifying lakes should be divided into “medium deep stratifying” and “deep stratifying” lakes to discern the naturally eutrophic and oligotrophic type.
3. The scientific community should consider to add descriptors or replace size and depth criteria by two alternative descriptors, which capture dominant biogeochemical issues related to morphometry: the share of epilimnion area water in contact with the sediment (by definition 100% in non-stratifying lakes) and the lake volume: watershed area ratio.
4. Reference lakes and reservoirs should not only be non-impacted (or minimally impacted) but should also be representative of physical conditions of their type. If salinity, turbidity, retention time etc. do not justify the establishment of separate types because of the small numbers of systems found, atypical lakes and reservoirs must not be selected as reference sites.
5. Despite the high number of indices developed during WFD implementation there is a need to critically review and further develop biological indices with the primary goal of eliminating ecologically erroneous species/genera/group assignments and the secondary goal of achieving greater unification. Future index development should be based both on structural and functional attributes of the community such as biodiversity and food web metrics.
6. Reference values of the key metric of phytoplankton method, the biovolume, should be ecologically sound to achieve comparability across Europe. Cyanobacteria biovolume used as a taxonomic or bloom metric should be the sum of all cyanobacterial species biovolume.
7. In accordance with the precautionary principle, special scrutiny is needed when class boundaries (e.g. in typology, reference conditions) set by one MS are conspicuously relaxed compared to other MSs.
8. Ecological expertise in the Universities should not be avoided in favor of consultancy contracts. Universities should function as supporting competent authorities.

## ACKNOWLEDGMENTS

We thank the Directorate for the Protection and Management of Water Resources of the Special Secretariat for Waters, Ministry of Environment and Energy for providing to M. M-G, member of the Greek National Committee for ecological classification, the lake data sets, the original physical-chemical and phytoplankton data (2012-2015) from the Greek National Water Monitoring Network.

